# *In vitro* Activity of Citrus IntegroPectin against Breast Cancer and Colon Cancer

**DOI:** 10.1101/2025.04.11.648349

**Authors:** Claudia D’Anna, Caterina Di Sano, Giovanna Li Petri, Giuseppe Angellotti, Rosaria Ciriminna, Mario Pagliaro

**Affiliations:** Istituto di Farmacologia Traslazionale, CNR, via U. La Malfa 153, 90146 Palermo, Italy; Istituto per lo Studio dei Materiali Nanostrutturati, CNR, via U. La Malfa 153, 90146 Palermo, Italy

**Keywords:** IntegroPectin, breast cancer, colon cancer, citrus, flavonoids

## Abstract

*In vitro* investigations of citrus IntegroPectin bioconjugates obtained through acoustic cavitation in water of different citrus fruit (lemon, red orange, and sweet orange) processing waste revealed the conjugate’s ability to inhibit the migratory features of Caco-2 colon cancer and MCF-7 breast cancer cells. All tested pectins substantially reduced cell migration of both tumor cell lines already at 0.5 and 1.0 mg/mL concentration. Dissolved in aqueous phase, furthermore, red orange and lemon IntegroPectin phytocomplexes at 5 and 10 mg/mL load reduced cell viability for both cell lines. Adding to promising results against lung cancer, these outcomes support further investigation of IntegroPectin citrus pectins for the treatment of cancer.

## 1. Introduction

Abundant in fruits and plants, pectin is a galacturonic acid polymer comprising homogalacturonan (HG), rhamnogalacturonan-I (RG-I), rhamnogalacturonan-II (RG-II), arabinogalacturonan, and xylogalacturonan regions.^[1]^ Its primary structure consists of repeating (1 → 4)-*α* -D-GalA (galactopyranosyluronic acid residues, partly methyl-esterified at *O*-6 position (and to a lower extent also acetyl-esterified at *O*-2 or *O*-3), interrupted by branched regions composed of (1 → 2)-*α* -l-rhamnose units (RG-I regions) further binding neutral sugars including galactose, arabinose, xylose, and fructose. Knowledge of its secondary structure, particularly in water, is expanding.^[2]^

Commercially sourced as high methoxyl (HM) pectin (degree of methyl esterification DE>50%) from citrus peels (or apple pomace) via prolonged hydrolysis using dilute mineral acid at 70–80 °C, pectin is purified via vacuum evaporation and alcohol precipitation using isopropyl alcohol to precipitate pectin.^[3]^ Low methoxyl (LM) pectin having DE < 50% is commercially produced by pectin manufacturers by controlled hydrolysis of HM pectin.

Pectin is a highly bioactive and health-beneficial substance.^[4]^ Its “modified” version rich in galactose residues present in RG-I galactan and arabinogalactan side chains obtained by hydrolysis at high pH of citrus pectin at 123°C for 1 h under 1.5 atm pressure (process affords lower molecular weight pectin by β-elimination and reduction in DE) binds to lung cancer cells galactoside-binding galactin-3 protein.^[5]^ Subsequent research found that said modified citrus pectin (MCP) is an anti-metastatic agent capable of inhibiting proliferation and metastasis for numerous cancers.^[6]^

Extracted via hydrodynamic^[7]^ (HC) or acoustic^[8]^ cavitation (AC) in water only of fresh industrial citrus processing waste (CPW) of different organically grown citrus fruits (lemon, orange, grapefruit and mandarin) juices sourced from, IntegroPectin is a LM pectin showing antioxidant, anti-inflammatory, cardioprotective, neuroprotective, mitoprotective, antimicrobial and anticancer properties.^[9]^ The broad-scope bioactivity of various IntegroPectin bioconjugates are ascribed to unique molecular structure (ultralow degree of methylation and abun-dant RG-I regions), enriched in citrus flavonoids (and terpenes).^[9]^

The vivid colors of lemon, red orange, and sweet orange IntegroPectin sourced via AC from fresh CPW (Fig.1) show visual evidence of the abundant amount of flavonoids contained in these bioconjugates. Using energy made available during the cavitation process, flavonoids abundant in citrus processing waste (chiefly present in the peel and residual pulp) can overcome the relatively low energy barrier and chemically bind to the galacturonic acid residues of pectin. Recently shown via a computational study,^[10]^ this explains the unique abundance of phenolics concentrated at the surface of citrus IntegroPectin.

**Figure 1.**
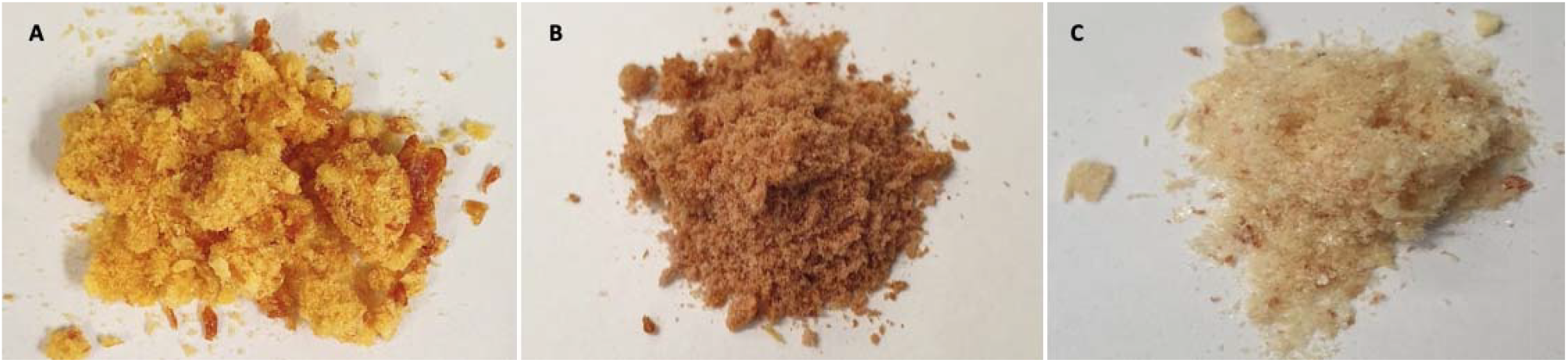
Sweet orange (A), red orange (B), and lemon (C) IntegroPectin obtained by acoustic cavitation.

Recently, we reported the *in vitro* activity against lung cancer of three citrus IntegroPectin bioconjugates sourced via AC of fresh lemon, red orange and sweet orange industrial processing waste evaluating both cell proliferation and long-term proliferation.^[11]^ Long-term proliferation and cell migration indeed are crucial processes in cancer progression. Metastasis arises from the ability of cancer cells to move from their original site to other parts of the body, forming secondary tumors, leading to uncontrolled growth. The development of effective therapies requires to prevent metastatic spread.^[12]^

In this study we report the outcomes of *in vitro* investigation of the anti-cancer properties of the aforementioned three citrus IntegroPectin bioconjugates using Caco-2 colon cancer and MCF-7 breast cancer cells. Originally sourced from a human colon adenocarcinoma, the human intestinal Caco-2 cell line is frequently used in biomedical studies as a model of the intestinal barrier.^[13]^ MCF-7 is the most studied human breast cancer cell line in the world, and results from this cell line have had a fundamental impact upon breast cancer research.^[14]^

Frequent in females, breast cancer is a significant public health problem that requires further research at the molecular level in order to define its prognosis and specific treatment.^[15]^ Similarly, colon cancer is a major cause of mortality in industrialized countries.^[16]^ Amid natural products researched as new therapies for both types of cancers, flavonoids are highly promising.^[17]^

## 2. Results

### 2.1 Citrus IntegroPectin bioconjugates

Recently reported,^[11]^ the outcomes of the analyses of selected flavonoids and phenolic acids in the three IntegroPectin (naringin, kaempferol, hesperidin, gallic acid, and p-coumaric acid) indicate that red orange IntegroPectin contains the highest concentration of the analyzed flavonoids and phenolic acids, with hesperidin exceeding the 7.2 mg/g concentration, and kaempferol amounting to nearly 0.12 mg/g. Sweet orange IntegroPectin also contains plentiful hesperidin, beyond 6.8 mg/g, an nearly 1.0 mg/g of naringin.

In agreement with the fact that hesperidin is by far the most abundant flavonoid in the peel of orange fruits, being particularly abundant in red orange cultivars.^[18]^ Lemon IntegroPectin contains nearly 3.5 mg/g of hesperidin but is the only IntegroPectin containing a substantial (2.6 mg/g) of p-coumaric acid, and nearly 2 mg/g (1.88 mg/g) of naringin.

The values of total phenolic content (TPC) assessed by the Folin-Ciocalteu method, expressed as milligrams of gallic acid equivalents (GAE) per gram of IntegroPectin, are exceptionally high for all three IntegroPectin phytocomplexes (33.6 mg GAE/mg for lemon, 30.91 mg GAE/mg for red orange, and 24.16 for sweet orange bioconjugates). For comparison, the TPC of lemon IntegroPectin sourced via HC and similarly dried via freeze-drying analyzed two years after its isolation is 0.88 mg GAE/g,^[19]^ whereas the TPC of lemon peel varies, depending on the cultivar, between 5.12 × 10^-3^ and 8.30 × 10^-3^ mg GAE/g).^[20]^

The radical scavenging activity of these bioconjugates assessed by the 2,2-diphenyl-1-picrylhydrazyl free radical (DPPH) assay shows pronounced and unique activity for all three IntegroPectin pectic materials. Indeed, as in the case of lemon and grapefruit IntegroPectin sourced via hydrodynamic cavitation,^[21]^ the antioxidant power of all the IntegroPectin bioconjugates, showing steep curves rather than a curve reaching a plateau as it happens for single flavonoids, indicates into an antioxidant power that is growing with time.

The XRD spectra (Fig.2) for all citrus IntegroPectin bioconjugates between 5° and 30° diffraction angle (2θ) displaying a broad peak centered around 18.5° show evidence that all citrus IntegroPectin phytocomplexes obtained via AC of citrus processing waste are comprised of amorphous pectin polymer.

**Figure 2.**
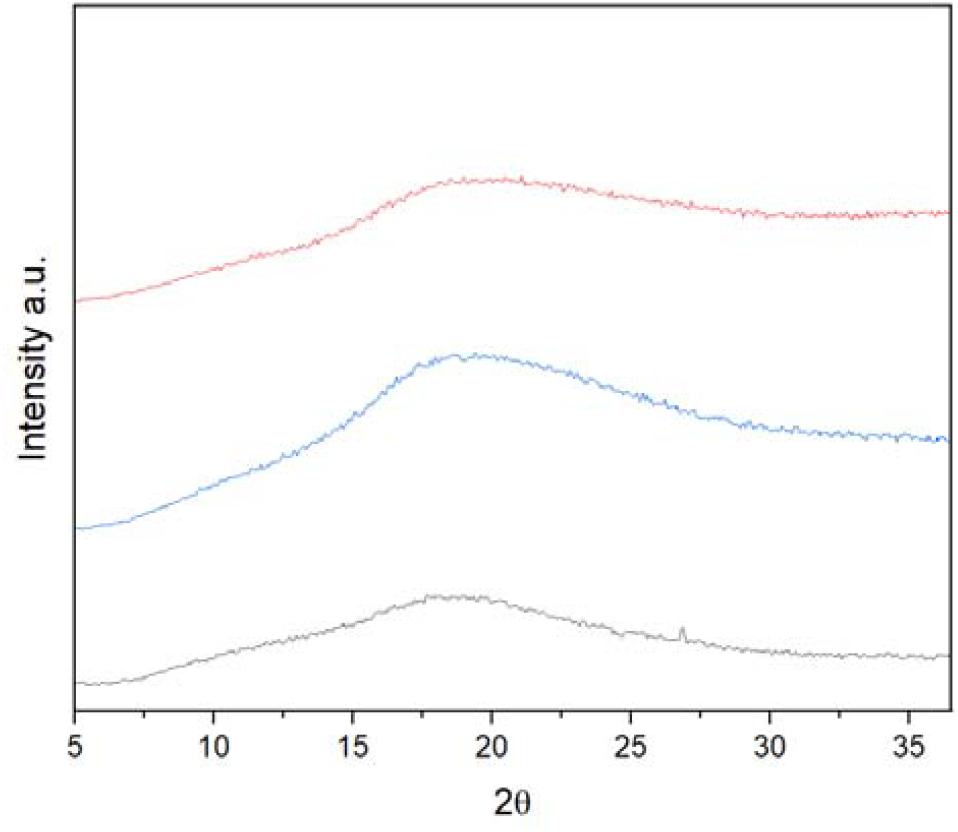
XRD spectra of lemon (blue line), sweet orange (red line), and red orange IntegroPectin (black line).

This indicates complete decrystallization of the HG regions, as it happens for lemon and grapefruit IntegroPectin sourced via HC. In either case (HC or AC), cavitation destroys the “fringed-micellar” structure of the crystalline regions of the semicrys-talline pectin biopolymer.^[22]^ For comparison, the XRD spectrum of commercial citrus pectin sourced via acid hydrolysis of citrus peel shows many diffraction peaks between 12.4° and 40.2° due to partly crystalline arrangement of the HG chains.^[23]^

The FTIR spectra of all IntegroPectin bioconjugates (Fig.3) are similar and clearly indicate highly de-esterified pectins rich in citrus flavonoids and phenolic acids.^[24]^ Signals at 1720 cm?^1^ and 1630 cm? ^1^ correspond to the stretching vibrations of the carbonyl group of esters and to carboxylate group of the pectin HG chain, respectively.^[24]^ Notably, the ratio between these two peaks varies among the IntegroPectin samples. This variation is particularly evident when comparing the spectra of lemon and red orange bioconjugate samples. In the case of lemon IntegroPectin, the two peaks exhibit similar intensities, whereas in the red orange spectrum, the signal at 1720 cm^-1^ is barely visible.

**Figure 3.**
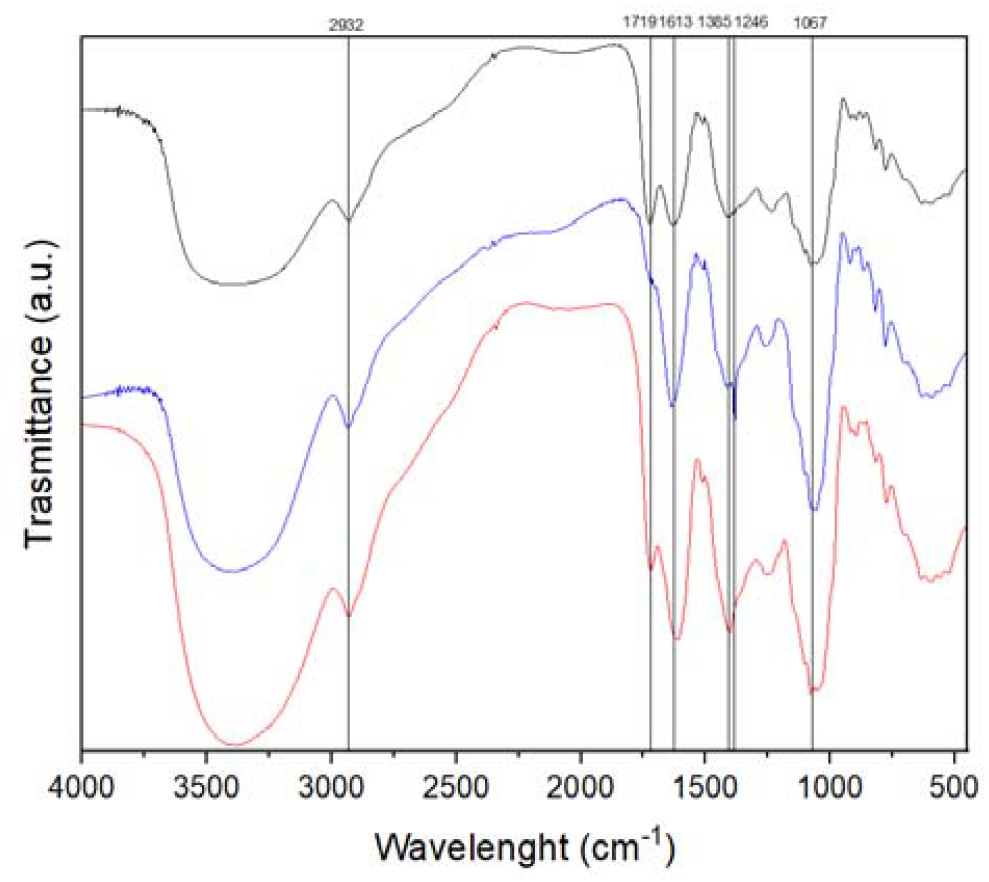
FTIR spectra of lemon (black line), red orange (red line), and sweet orange (blue line) IntegroPectin.

This difference can be attributed to the degree of esterification, with the red orange sample showing the lowest DE. Finally, the signal at 2931 cm^−1^ is due to the stretching vibrations of C–H bonds of CH and CH_2_ groups of polysaccharide rings, whereas the broad band centered at 3406 cm^−1^ is due to the O– H stretching vibration of the pyranose ring and adsorbed water.

### 2.2 Effects of IntegroPectin from lemon, red orange and sweet orange on Caco-2 cell viability

Fig.4 shows evidence that the cell viability of Caco-2 cells cultured with citrus IntegroPectin bioconjugaes in aqueous solution at increasing concentration (0.25, 0.5, 1.0, 5.0 and 10 mg/mL) for 24 h started to decrease at 5 mg/mL IntegroPectin concentration.

**Figure 4.**
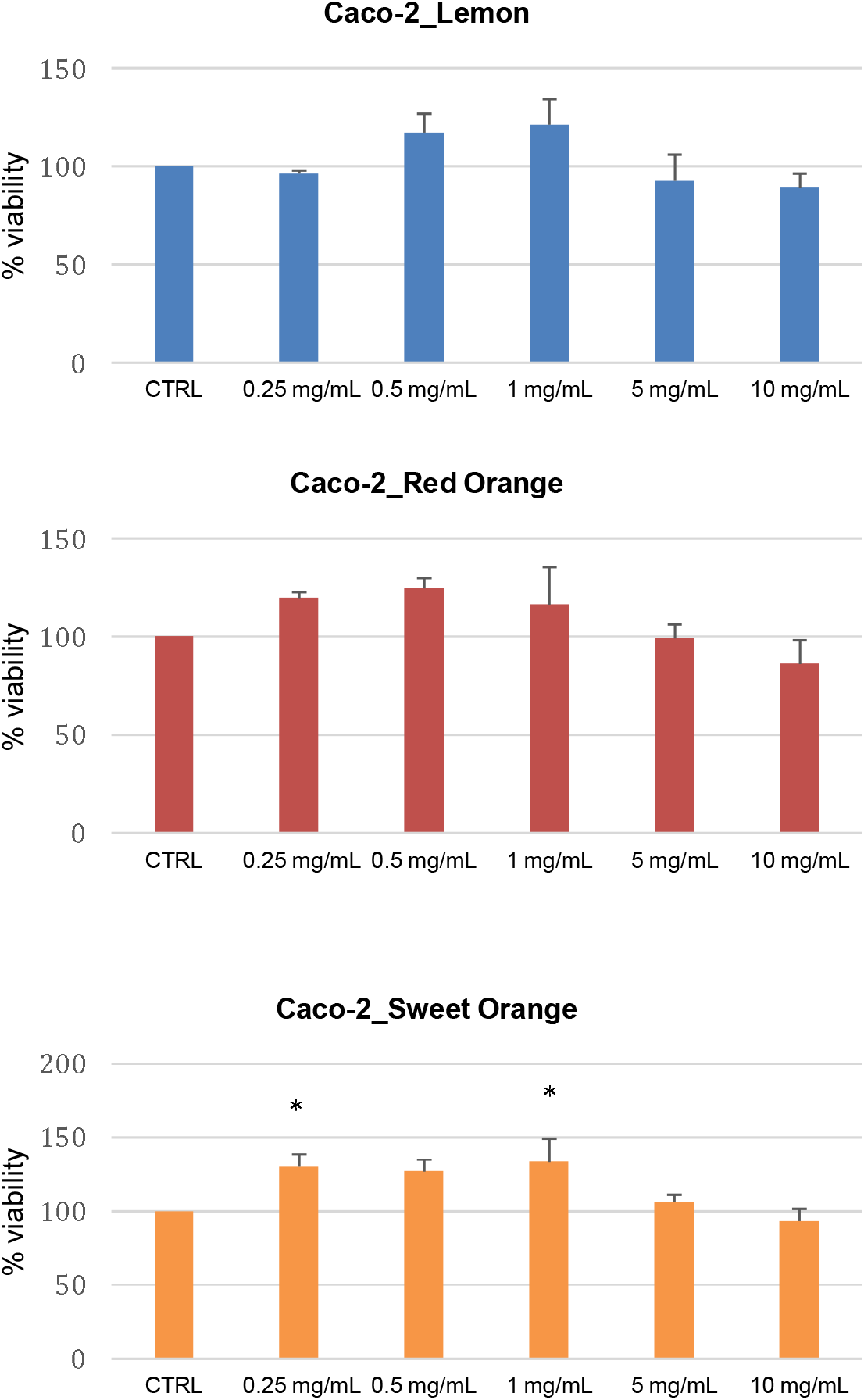
Effect of different citrus IntegroPectin bioconjugates on Caco-2 cell viability. Cells were cultured for 24 h with different citrus IntegroPectin bioconjugates (0.25, 0.5, 1.0, 5.0 and 10 mg/mL). Data are expressed as % of untreated and represent mean ± SD (*n*=3).

Cell viability was measured through the MTS cell viability test, in which reduction of MTS (3-(4,5-dimethylthiazol-2-yl)-5-(3-carboxymethoxyphenyl)-2-(4-sulfophenyl)-2H-tetrazolium salt) is due to dehydrogenases enzymes present in mitochondria, with this endpoint being a good biomarker of the damage induced in this organelle. Added to the medium, MTS is reduced by cells into colored formazan which is measured spectrophotometrically at 490 nm after 30 min of incubation in the dark.^[25]^

In the case of lemon and red orange bioconjugate a concentration of 5 mg/ml reduced cell viability below 90%. The lowest cell viability was observed for red orange IntegroPectin at 10 mg/mL concentration, when Caco-2 cell viability dropped to 86.3%. The IntegroPectin sourced from sweet orange was the least active in reducing cell viability, that reached 93.4% at 10 mg/mL load but increased beyond 100% for all loads comprised between 0.25 and 5 mg/mL.

### 2.3. Effects of IntegroPectin from lemon, red orange and sweet orange on MCF-7 cell viability

Plots in Fig.5 show that a significant reduction in MCF-7 cell colony number was observed after treatment of the cells for 24 h with red orange IntegroPectin at 10 mg/mL concentration when cell viability was 88.2%. Viability reduction below 100% (94.1%) was observed already at 5 mg/mL concentration.

**Figure 5.**
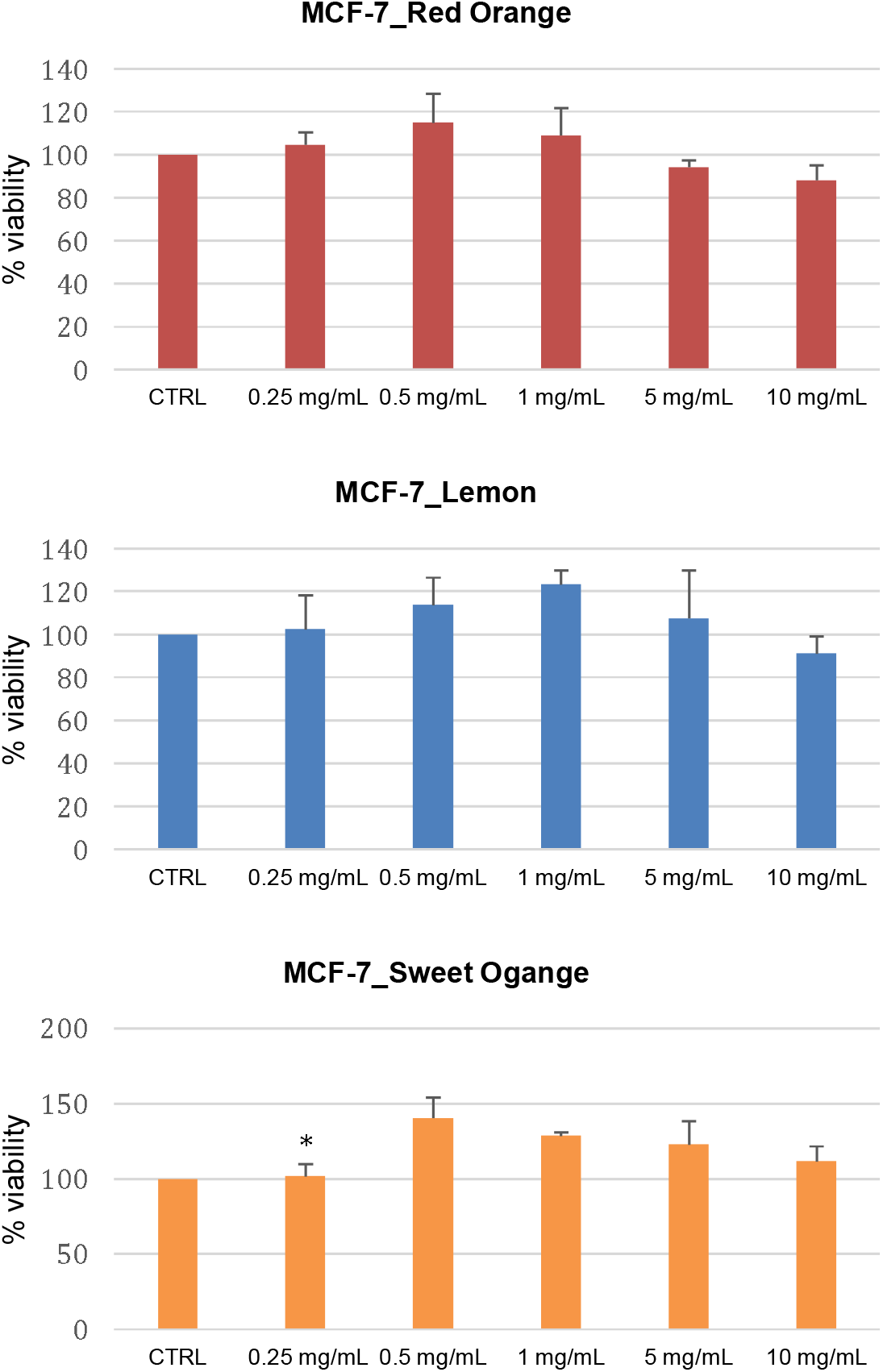
Effect of different citrus IntegroPectin bioconjugates on MCG-7 cell viability. Cells were cultured for 24 h with different citrus IntegroPectin bioconjugates (0.25, 0.5, 1.0, 5.0 and 10 mg/mL). Data are expressed as % of untreated and represent mean ± SD (*n*=3).

Similarly, in the case of lemon IntegroPectin, reduction of cell viability to 91% was observed at 10 mg/mL concentration. In the case of sweet orange IntegroPectin, on the other hand, cell viability increased at all tested loads.

Concerning the cell shape changed in the presence of IntegroPectin bioconjugates in solution. Photographs in Fig.6 and Fig.7 show that whereas at loads between 0.25 and 1.0 mg/mL the morphology of both Caco-2 and MCF-7 cells was not significantly affected, at higher loads tested of 5 and 10 mg/mL, the cell shape significantly changed for both cell lines.

**Figure 6.**
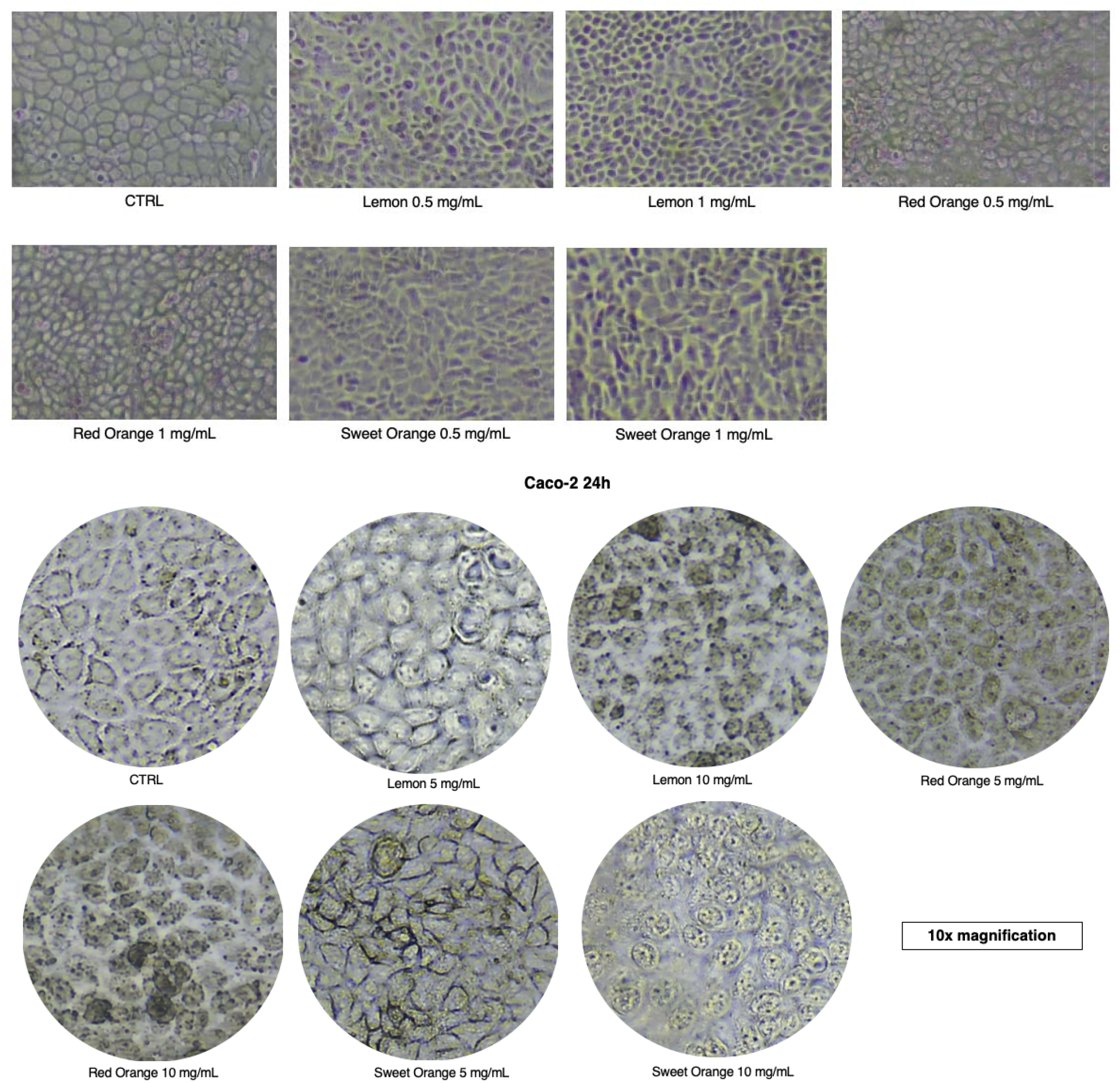
Morphology of Caco-2 cells in the presence of different citrus IntegroPectin bioconjugates at 0.5 and 1.0 mg/mL load after 24 h (*top*); and in the presence of the same IntegroPectin bioconjugates at 5 and 10 mg/mL load (*bottom*).

**Figure 7.**
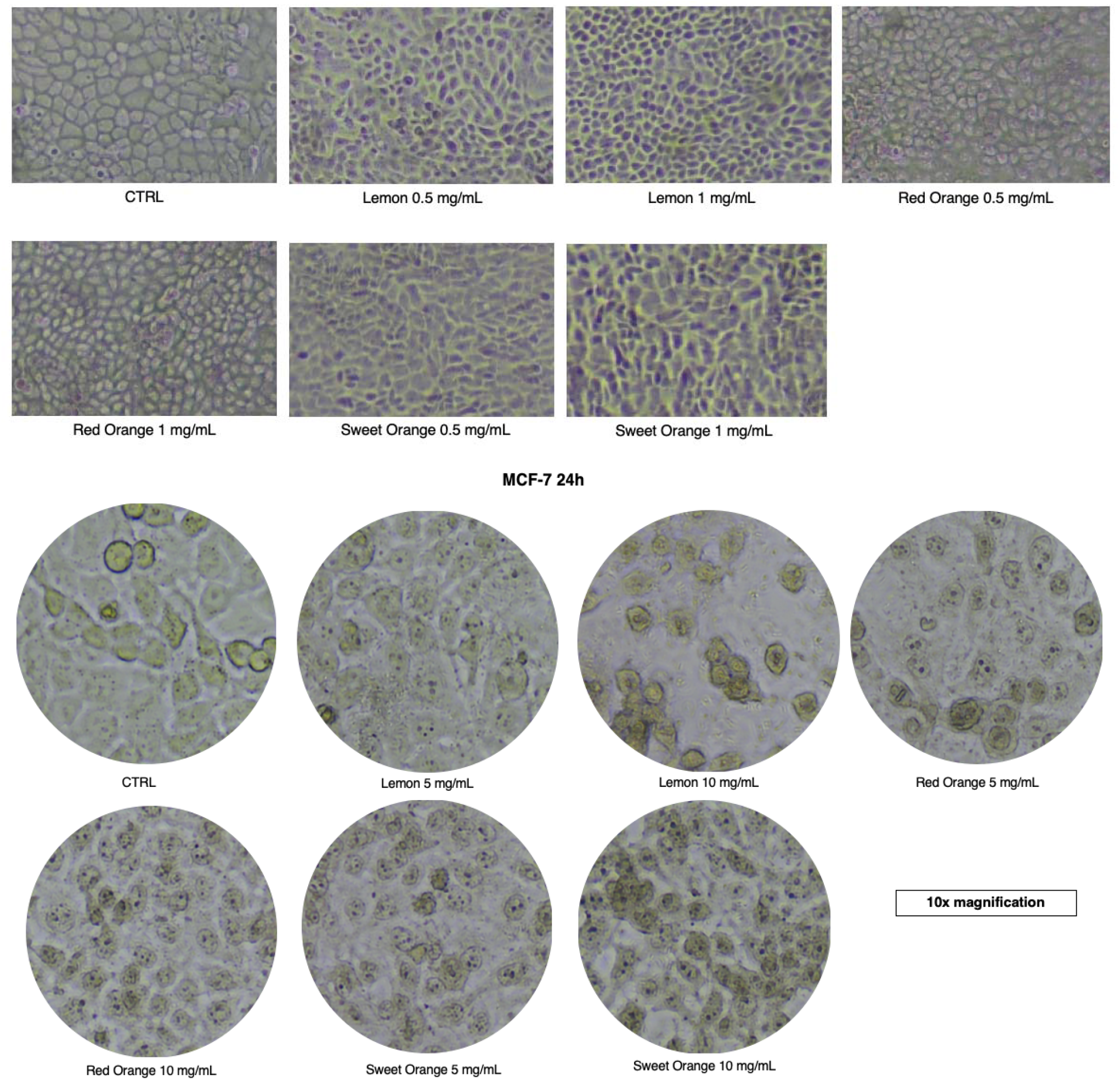
Morphology of MCF-7 cells in the presence of different citrus IntegroPectin bioconjugates at 0.5 and 1.0 mg/mL load after 24 h (top), and in the presence of the same IntegroPectin at 5 and 10 mg/mL load (bottom).

In the case of Caco-2 cells in the presence all (lemon, red orange, and sweet orange) IntegroPectin phytocomplexes at 10 mg/mL load, cells darkened, detached from each other and contracted pointing to changes in the structure of the cholesterol-rich membrane and thus to endocytosis. It is relevant here to notice that in Caco-2 cells in contact with curcumin-loaded zein/pectin nanoparticles endocytosis takes place via caveolae-dependent endocytosis, clathrin-dependent endocytosis, and micropinocytosis, indicating that nanoparticle size, composition and hydrophilicity/hydrophobicity all affect the nanoparticle internalization and transport across the cells.^[26]^

### 2.4. Effects of IntegroPectin from lemon, red orange and sweet orange on Caco-2 cell line motility

Increased cancer cell motility is a feature of tumor aggressiveness.^[27]^ Photographs and histograms in Fig.8 show that Caco-2 cells stimulated with different citrus IntegroPectin bioconjugates at 0.5 and 1 mg/mL concentration for 24 and 48 h significantly reduced cell motility.

**Figure 8.**
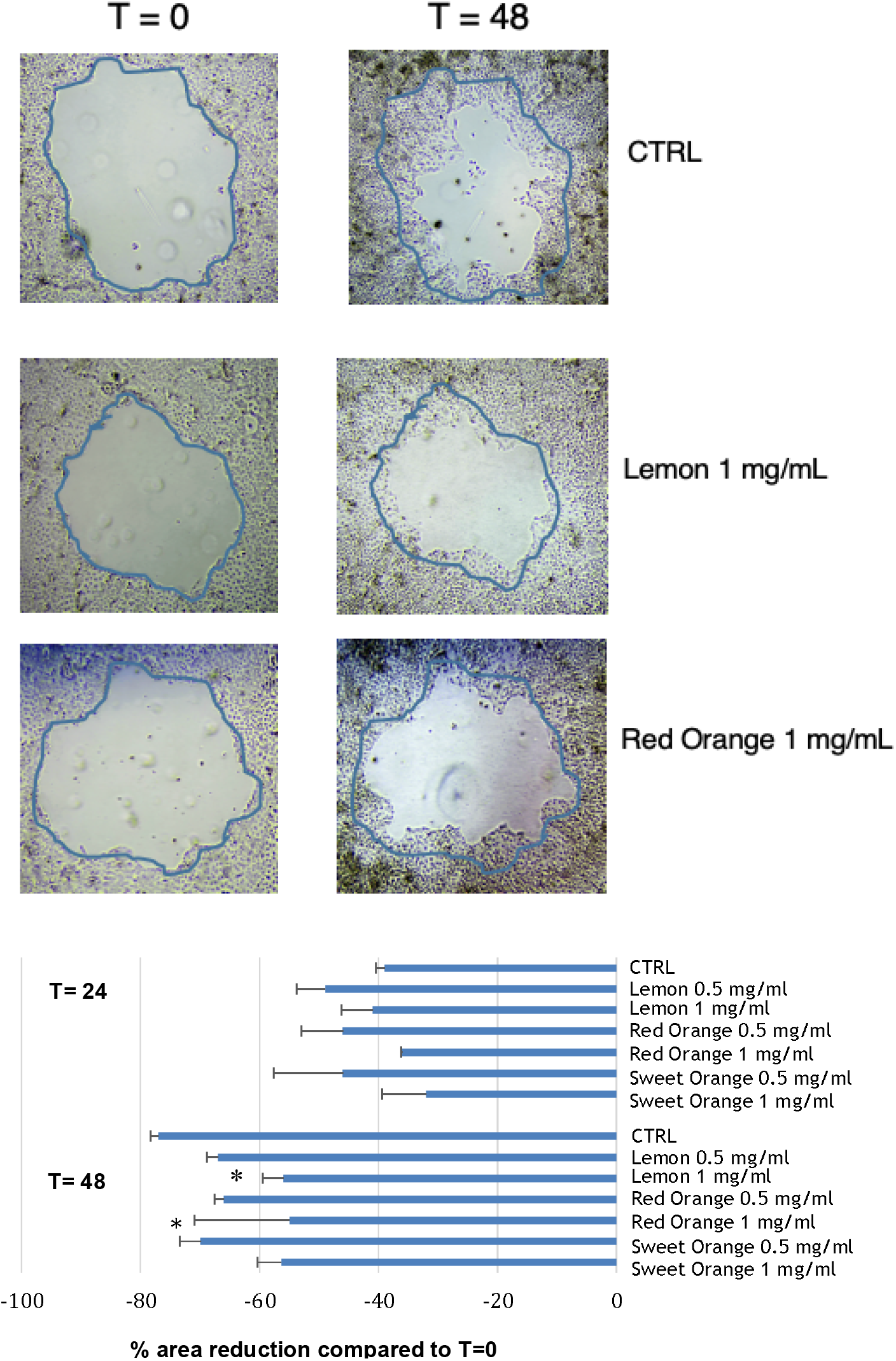
Effect of different citrus IntegroPectin bioconjugates on cell migration, in Caco-2 cell line. Cells were stimulated with different Citrus IntegroPectin bioconjugates at 0.5 and 1 mg/mL concentration, for 24 and 48 h.

The most effective were lemon and red orange IntegroPectin conjugates that at 1.0 mg/mL concentration that after 48 h limited the wound area reduction to just 55% vs. 77% of the untreated cells. Contrary to what observed for cell viability in the MTS test, the sweet orange IntegroPectin too significantly reduced cell migration, with 56% wound area reduction after 48 h vs. 77% in the case of untreated cells. This outcome is in agreement with the previously reported ability of sweet orange IntegroPectin bioconjugate to reduce cell motility also in the case of A549 lung cancer cells.^[11]^

As shown by photographs and plots in Fig.9, in the case of MCF-7cells too, all three IntegroPectin bioconjugates were able to significantly decrease cell migration. Now, the most effective citrus bioconjugates were red orange and sweet orange IntegroPectin, that at 1.0 mg/mL concentration after 48 h respectively limited the wound area reduction to just 40.2% *vs*. 75.9% of the untreated cells, and to 42.2% in the case of sweet orange IntegroPectin.

**Figure 9.**
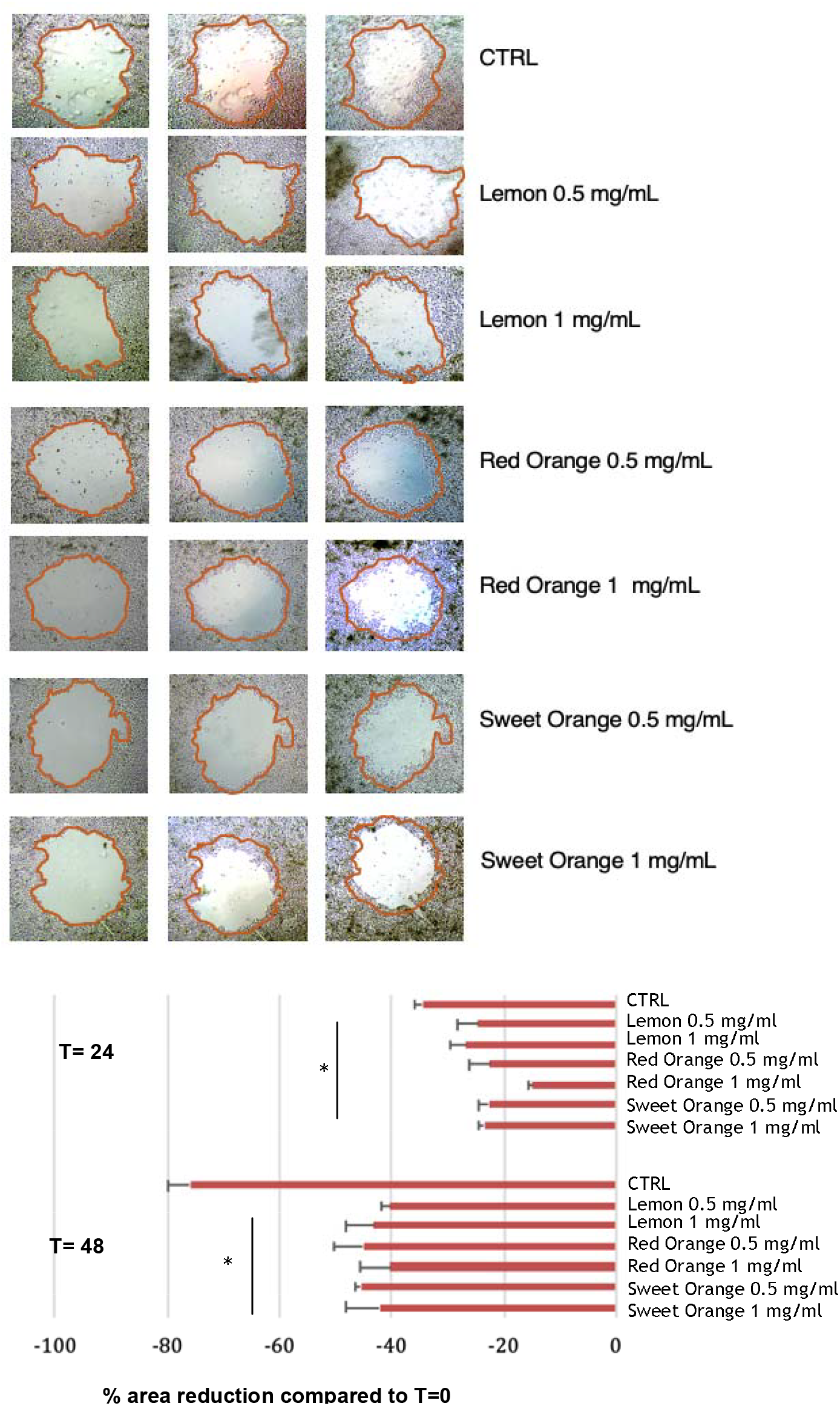
Effect of different citrus IntegroPectin bioconjugates on cell migration, in MCF-7cell line. Cells were stimulated with different citrus IntegroPectin bioconjugates at 0.5 and 1 mg/mL concentration, for 24 and 48 h.

Lemon IntegroPectin at 1.0 mg/mL concentration too after 48 h limited the wound area reduction to 43.2%, but in this case even more pronounced limited wound area reduction to 40.3% was observed at 0.5 mg/mL concentration. Due to enhanced ability to detach and plasticity driven by epithelialmesenchymal transition (transformation, and metastasis of cancer cells through blood vessels and lymphatics) which endows the cells with increased invasiveness and motility),^[28]^ tumor cell circulation is a key step in the metastatic process.

## 3. Discussion

Following demonstration of antiproliferative activity *in vitro* of IntegroPectin bioconjugates sourced via AC of lemon and red orange industrial processing waste (and isolated via freeze-drying) against lung cancer cells,^[10]^ these findings demonstrate that these new citrus pectins possess significant anticancer activity *in vitro* against MCF-7 breast cancer and Caco-2 colon cancer cells, inhibiting both proliferation and cell motility.

Once again this bioactivity may be due to the unique molecular structure of these new citrus pectins having low degree of esterification, enriched in RG-I regions, and rich in molecularly bound citrus flavonoids (and adsorbed terpenes).^[9]^

Accordingly, the significant improvement in biological activity and bioavailability of MCP have both been ascribed to its low DE and low molecular weight, translating into higher number of free carboxylic acid groups binding and inhibiting galactin-3.^[29]^

These structural features suggest a suitable mechanism explaining the broad-scope anticancer activity *in vitro* of citrus IntegroPectin bioconjugates. Alongside possessing the intrinsic anticancer activity of citrus pectin rich in RG-I regions and free carboxylic acids, the highly soluble IntegroPectin bioconjugates uniquely rich in molecularly bound flavonoids deliver otherwise poorly soluble citrus flavonoids such as hesperidin, naringin, kaempferol and eriocitrin within the cell membranes, where these flavonoids can ultimately unleash their anticancer activity so far limited by their lack of solubility in water. Indeed, the poor oral bioavailability of these and other dietary biophenols has limited for decades their development as therapeutic agents.^[30,31]^ For example, the *in vivo* plasma concentrations of flavonoids are typically in the range of 0.01–0.1 μM, significantly lower than the half maximal inhibitory concentration or half maximal effective concentration) values of 5–50 μM commonly reported for their anticancer, anti-inflammatory effects and other effects *in vitro*.^[32]^

Indeed, studying the cell viability of Caco-2 cells treated with 40 μM flavonoid, Xu and co-workers found no significant difference between the Caco-2 cell viability of the control and each flavonoid, showing that flavonoids (including citrus flavonoids such as quercetin, rutin, naringenin, kaempferol and hesperetin) at the tested concentration showed no cytotoxicity to Caco-2 cells.^[33]^ Indeed, when a quercetin-DHA ester derivative (DHA is docosahexaenoic acid) is covalently bound to pectin, the conjugate becomes able to inhibit migration and cell viability of Caco-2 cells.^[34]^

This approach aimed at improving the bioavailability of flavonoids via covalent binding to pectin is general. It has for example been used also to enhance the bioavailability of naringin with the resulting conjugate showing significantly enhanced, antioxidant, antimicrobial and anti-cancer activity.^[35]^ The outcome was ascribed to enhanced delivery of naringenin bound to water-soluble pectin, making pectin a “versatile and effective means of transporting polyphenol compounds”.^[36]^

On the other hand, the fact that sweet orange IntegroPectin even increases Caco-2 cell viability up to a concentration of 1 mg/mL (Fig.2) is in agreement with the fact that the viability of Caco-2 cells incubated with different hesperidin concentrations ranging from 0 to 100□μM remained above 90% compared with that of untreated control cells, namely cell viability values considered non-cytotoxic.^[37]^

## 4. Conclusions

In summary, we have discovered that citrus IntegroPectin bioconjugates obtained from lemon and red orange industrial processing waste through acoustic cavitation conducted in water only at room temperature followed by IntegroPectin isolation via freeze drying show substantial anticancer action *in vitro* against human colon (Caco-2) and breast (MCF-7) cancer cells.

Red orange and lemon IntegroPectin phytocomplexes affected long-term proliferation and cell migration, whereas sweet orange did reduce (but rather increased) colony formation activity. These results indicate that citrus IntegroPectin bioconjugates exhibit antitumor effects these cell lines by reducing cancer cell progression, providing a basis for further investigation in lung cancer therapy. The use of IntegroPectin provides a new approach, alternative to *in vitro* and *in vivo* studies on the anti-cancer effects of citrus pectin conducted using either commercial citrus or citrus pectin sourced from the fruit peels using conventional acid hydrolysis at 70-80 °C.^[38]^

Adding to the relevance of said *in vitro* results is the fact that citrus pectin is a health beneficial dietary component ubiquitous in fruit and plants. Present in modest amounts in citrus fruit (orange, lemon and grapefruit) juice, citrus flavonoids too are highly beneficial for human health. As mentioned above, their biomedical applications so far have been limited by ultralow solubility in water.^[30,31]^

Cavitation of citrus processing waste achieves in one-step and in one-pot the concomitant extraction and structural modification of pectin, converted in pectin enriched in RG-I regions, having ultra-low DE and containing an exceptional amount of flavonoids bound to the pectin backbone. Remarkably, Zheng and co-workers lately first confirmed that hesperidin molecularly bonds to pectin in citrus peel, as anticipated in a computational study; and that citrus flavonoids including hesperidin can be enzymatically conjugated to citrus pectin at high rate (20.21%) affording a bioconjugate showing better emulsifying properties.^[39]^

Commercial citrus pectin does not show anticancer activity against breast cancer cells. However, pointing once again to the crucial relevance of citrus pectin’s structure, heat-modified citrus pectin (MCP) at 3 mg/mL concentration inhibits breast cancer development in mice by targeting tumorassociated macrophage survival and polarization in hypoxic microenvironment.^[40]^ Similarly, a number of preclinical studies have shown that kaempferol inhibits the growth, induces apoptosis of, and inhibits the migration and invasion of breast cancer cells.^[41]^ Yet, poor bioavailability requires the development of drug delivery systems capable to enhance delivery and thus activity of this flavonoid in biomedically relevant conditions.^[41]^

The process to produce citrus IntegroPectin, via cavitation of citrus processing waste originating from citrus fruit organically farmed, is both reproducible and readily scalable.^[8]^ Colon and breast cancers are public health problems. Hence, preclinical and clinical studies to test the activity of citrus IntegroPectin bioconjugates in the treatment of these (and other) cancers should be urgently conducted.

## Experimental section

### IntegroPectin isolation, flavonoid quantification, DPPH assay, and Folin–Ciocalteu assay

Lemon, red orange, and sweet orange IntegroPectin bioconjugates were prepared via AC of fresh citrus processing waste kindly provided by Citrus Campisi (Siracusa, Italy) as recently reported.^[11]^ The same study included details on flavonoid quantification, DPPH assay, and Folin–Ciocalteu assay.

### IntegroPectin solutions in PBS

A sample of each IntegroPectin was dispersed in PBS (phosphate-buffered saline, pH 7.4, purchased from Gibco Invitrogen, New York, USA) at a concentration of 20 mg/mL. The resulting mixture was sonicated for 3 min to achieve a homogeneous solution. For each IntegroPectin, the resulting solution was stored at 4 °C prior to conduct the biological tests

### Cell Cultures

MCF-7 and Caco-2 cells were cultured in D-MEM and RPMI medium, respectively. Both medium were supplemented with heat-deactivated (56°C, 30□min) 10% FBS, streptomycin and penicillin, 1% nonessential amino acids and 2□mM L-glutamine (all purchased from Euroclone, Pero, Italy). The cells grown as adherent monolayers, and reached approximately 80% of confluence, they were treated with IntegroPectin from lemon red orange and sweet orange at concentrations of 0.5 and 1 mg/mL and incubated for 24 h in a humidified ambient (5% air CO_2_) at 37 °C.

### Effects of IntegroPectin from lemon, red orange and sweet orange on Caco-2 and MCF-7 cell viability

#### MTS

Cell viability was assessed with the CellTiter 96 Aqueous One Solution Cell Proliferation Assay (PROMEGA, Madison WI, USA), in accordance with the manufacturer’s guidelines. Cells were cultured in 96-well plates and treated for 24 h in quadruplicate with IntegroPectin from lemon red orange and sweet orange at different concentrations (0.25, 0.5, 1.0, 5.0 and 10 mg/mL). Absorbance was measured at 490 nm using a microplate reader. The results were expressed as a percentage relative to the untreated (NT) control and the data are presented as the average of quadruplicate wells ± standard deviation (SD).

### Effect of IntegroPectin from lemon, red orange and sweet orange on A549 cell line motility

During the experiment, a “wound” (a cell-free zone in the cell monolayer) is made and recolonization of the scratched region is monitored to quantify cell migration area. In detail, the wound area is tracked and the ratio between wound area at time *t* A(*t*) and the initial area A(0) expressed as the percentage wound area at a specific time point indirectly allows to evaluate the migration rate.^[42]^

Briefly, MCF-7 and Caco-2 cells were grown in a 6-well plate until they reached confluence. Three circular wounds were created in each well using a 200-μl pipette tip. Wells were washed with PBS to discharge any debris, and after 1 h, IntegroPectin from lemon, red orange and sweet orange added (0.5 and 1 mg/mL concentrations). Images were performed at 0, 24, and 48 h post-wounding, with a digital camera integrated into an inverted phase-contrast microscope.

To analyze wound area and closure rate, we used the ImageJ software, presenting results as percentage of area reduction at 24 h and 48 h, relative to time zero (0 h). Differences between the various experimental conditions were assessed using ANOVA, with Fisher’s test applied for corrections (*p < 0.05 was considered statistically significant) using free software for analysing numbers and sizes of cell colonies.^[43]^

## Acknowledgements

We thank Citrus Campisi (Siracusa, Italy) for the generous gift of industrial processing waste of organically grown citrus fruits from which the IntegroPectin bioconjugates were sourced. Work of G.L.P. was supported by MICS (Made in Italy - Circular and Sustainable) Extended Partnership and received funding from the European Union NextGenerationEU (PNRR - Mission 4 Component 2, Investment 1.3 - D.D.1551.11-10-2022, PE00000004). Work of G.A. was supported by the SAMOTHRACE (Sicilian Micro and Nano TechnologyResearch and Innovation Center) Innovation Ecosystem and received funding from European Union NextGenerationEU (PNRR – Mission 4 Component 2, Investment 1.5 (ECS00000022)). R.C. and M.P. thank Ministero dell’Universita e della Ricerca for funding, under Progetto “FutuRaw. Le materie prime del futuro da fonti non-critiche, residuali e rinnovabili”, Fondo Ordinario Enti di Ricerca, 2022, (CUP B53C23008390005).

## Data Availability Statement

The data that support the findings of this study are available from the corresponding authors upon reasonable request.

## References

[1] D. Ropartz, M. C. Ralet, Pectin Structure. In: Pectin: Technological and Physiological Properties, V. Kontogiorgos (Ed.), Springer, Cham: 2020; pp.17–36.

[2] A. Zdunek, P.M. Pieczywek, J. Cybulska, The primary, secondary and structures of higher levels of pectin polysaccharides. Compr. Rev. Food Sci. Food Saf. 2021, 20, 1101–1117. DOI:10.1111/1541-4337.12689

[3] Z.K. Muhidinov, I. Khurshed, I. Ikromi, et al., Structural characterization of pectin obtained by different purification methods. Int. J. Biol. Macromol. 2021, 183, 2227–2237. DOI:10.1016/j.ijbiomac.2021.05.094

[4] S. Zhang, G. I. N. Waterhouse, F. Xu, et al., Recent advances in utilization of pectins in biomedical applications: a review focusing on molecular structure-directing health-promoting properties. Crit. Rev. Food Sci. Nutr. 2021, 63, 3386–3419. DOI:10.1080/10408398.2021.1988897

[5] D. Platt, A. Raz, Modulation of the lung colonization of B16-F1 melanoma cells by citrus pectin. J. Natl. Cancer Inst. 1992, 84, 438–442. DOI:10.1093/jnci/84.6.438

[6] V. V. Glinsky, A. Raz, Modified citrus pectin anti-metastatic properties: one bullet, multiple targets. Carbohydr. Res. 2009, 344, 1788–1791. DOI:10.1016/j.carres.2008.08.038

[7] F. Meneguzzo, C. Brunetti, A. Fidalgo, et al., Real-scale integral valorization of waste orange peel via hydrodynamic cavitation. Processes 2019, 7, 581. DOI:10.3390/pr7090581

[8] R. Ciriminna, F. Meneguzzo, G. Li Petri, et al., Cavitation as a zero-waste circular economy process to convert citrus processing waste into biopolymers in high demand. J. Biores. Bioprod. 2024, 9, 246–252. DOI:10.1016/j.jobab.2024.09.002

[9] R. Ciriminna, V. Di Liberto, C. Valenza, et al., Citrus IntegroPectin: a multifunctional bioactive phytocomplex with large therapeutic potential. ChemRxiv 2024, DOI:10.26434/chemrxiv-2024-45v17

[10] V. Butera, R. Ciriminna, C. Valenza, et al., Citrus IntegroPectin: a computational insight. Discover Mol. 2025, 2, 6. DOI:10.1007/s44345-025-00013-z

[11] C. Di Sano, C. D’Anna, G. Li Petri, et al., In vitro activity of citrus IntegroPectin against lung cancer cells. bioRxiv 2025, 01.15.633201. DOI:10.1101/2025.01.15.633201

[12] R.L. Anderson, T. Balasas, J. Callaghan, et al., A framework for the development of effective anti-metastatic agents. Nat. Rev. Clin. Oncol. 2019, 16, 185–204. DOI:10.1038/s41571-018-0134-8

[13] Y. Sambuy, I. De Angelis, G. Ranaldi, et al. The Caco-2 cell line as a model of the intestinal barrier: influence of cell and culture-related factors on Caco-2 cell functional characteristics. Cell Biol. Toxicol. 2005, 21, 1–26. DOI:10.1007/s10565-005-0085-6

[14] (a) S. Comsa, A. M. Cimpean, M. Raica, The story of MCF-7 breast cancer cell line: 40 years of experience in research. Anticancer Res. 2015, 35, 3147–3154; (b) A. V. Lee, S. Oesterreich, N. E. Davidson, MCF-7 cells—changing the course of breast cancer research and care for 45 years. J. Natl. Cancer Inst. 2015, 107, djv073. DOI:10.1093/jnci/djv073

[15] X. Xiong, L.W. Zheng, Y. Ding, et al. Breast cancer: pathogenesis and treatments. Sig. Transduct. Target Ther. 2025, 10, 49. DOI:10.1038/s41392-024-02108-4

[16] R. L Siegel, A.N. Giaquinto, A. Jemal, Cancer statistics, 2024. CA Cancer J. Clin. 2024, 74, 12–49. DOI:10.3322/caac.21820

[17] A. Zaitun Hasibuan, Y. Simanjuntak, Ev. Hey-Hawkins, et al., Unlocking the potential of flavonoids: Natural solutions in the fight against colon cancer. Biomed. Pharmacother. 2024, 176, 116827. DOI:10.1016/j.biopha.2024.116827

[18] J. A. Manthey, K. Grohmann, Concentrations of hesperidin and other orange peel flavonoids in citrus processing byproducts. J. Agric. Food Chem. 1996, 44, 811–814. DOI:10.1021/jf950572g

[19] D. Nuzzo, L. Cristaldi, M. Sciortino, et al., Exceptional antioxidant, non-cytotoxic activity of integral lemon pectin from hydrodynamic cavitation. ChemistrySelect 2020, 5, 5066–5071. DOI:10.1002/slct.202000375

[20] W. Xi, J. Lu, J. Qun, et al., Characterization of phenolic profile and antioxidant capacity of different fruit part from lemon (Citrus limon Burm.) cultivars. J. Food Sci. Technol. 2017, 54, 1108–1118. DOI:10.1007/s13197-017-2544-5

[21] G. Di Prima, A. Scurria, G. Angellotti, et al., Grapefruit IntegroPectin isolation via spray drying and via freeze drying: A comparison. Sustain. Chem. Pharm. 2022, 29, 100816. DOI:10.1016/j.scp.2022.100816

[22] R.M. Gohil, Synergistic blends of natural polymers, pectin and sodium alginate. J. Appl. Polym. Sci. 2011, 120, 2324–2336. DOI:10.1002/app.33422

[23] W. Wang, X. Ma, P. Jiang, et al., Characterization of pectin from grapefruit peel: A comparison of ultrasound-assisted and conventional heating extractions. Food Hydrocol. 2016, 61, 730–739. DOI:10.1016/j.foodhyd.2016.06.019

[24] R. Ciriminna, A. Fidalgo, D. Carnaroglio, et al., Eco-friendly extraction of pectin and essential oils from orange and lemon peels. ACS Sustain. Chem. Eng. 2016, 4, 2243–2251. DOI:10.1021/acssuschemeng.5b01716

[25] J.A. Baltrop, T.C. Owen, A.H. Cory, et al., 5-((3-Carboxyphenyl)-3-(4,5-dimethylthiazolyl)-3-(4-sulfophenyl)) tetrazolium, inner salt (MTS) and related analogs of 2-(4,5-dimethylthiazolyl)-2,5-diphenylterazolium bromide (MTT) reducing to purple water soluble formazan as cell-viability indicators. Bioorg. Med. Chem. Lett. 1991, 1, 611. DOI:10.1016/S0960-894X(01)81162-8

[26] Y. Peng, X. Li, P. Gu, et al., Curcumin-loaded zein/pectin nanoparticles: Caco-2 cellular uptake and the effects on cell cycle arrest and apoptosis of human hepatoma cells (HepG2). J Drug Deliv. Sci. Technol. DOI:10.1016/j.jddst.2022.103497 2022, 74, 103497.

[27] X. Gu, S. Wei, X. Lv, Circulating tumor cells: from new biological insights to clinical practice. Sig. Transduct. Target Ther. 2024, 9, 226. DOI:10.1038/s41392-024-01938-6

[28] D. Nuzzo, M. Scordino, A. Scurria, et al., Protective, antioxidant and antiproliferative activity of grapefruit IntegroPectin on SH-SY5Y cells. Int. J. Mol. Sci. 2021, 22, 9368. DOI:10.3390/ijms22179368

[29] L. An, G. Chang, L. Zhang, et al. Pectin: Health-promoting properties as a natural galectin-3 inhibitor. Glycoconj. J. 2024, 41, 93–118. DOI:10.1007/s10719-024-10152-z

[30] A. Scalbert, G. Williamson, Dietary intake and bioavailability of polyphenols. J. Nutr. 2000, 130, 2073S–2085S. DOI:10.1093/jn/130.8.2073S

[31] S. Elmeligy, M. Hathout, S.A.M. Khalifa, et al., Pharmaceutical manipulation of citrus flavonoids towards improvement of its bioavailability and stability. A mini review and a meta-analysis study. Food Biosci. 2021, 44, 101428. DOI:10.1016/j.fbio.2021.101428

[32] P.A. Kroon, M.N. Clifford, A. Crozier, et al., How should we assess the effects of exposure to dietary polyphenols in vitro? Am. J. Clin. Nutr. 2004, 80, 15–21. DOI:10.1093/ajcn/80.1.15

[33] Y. Fang, W. Cao, M. Xia, et al., Study of structure and permeability relationship of flavonoids in Caco-2 cells. Nutrients 2017, 9, 1301. DOI:10.3390/nu9121301

[34] G. Carullo, U.G. Spizzirri, R. Malivindi, et al., Development of quercetin-DHA ester-based pectin conjugates as new functional supplement: effects on cell viability and migration. Nutraceuticals 2022, 2, 278–288. DOI:10.3390/nutraceuticals2040021

[35] J. Mundlia, M. Ahuja, P. Kumar, et al., Improved antioxidant, antimicrobial and anticancer activity of naringenin on conjugation with pectin. Biotech 2019, 9, 312. DOI:10.1007/s13205-019-1835-0

[36] V. Kumar R., D. Srivastava, A. Verma, et al., Biosynthesis of pectin and its vital role in the bioavailability of phytochemicals associated with therapeutic properties. Pharmacol. Res. Nat. Prod. 2025, 7, 100214. DOI:10.1016/j.prenap.2025.100214

[37] H.-Y. Park, J.-H. Yu, Hesperidin enhances intestinal barrier function in Caco-2 cell monolayers via AMPK-mediated tight junction-related proteins. FEBS Open Bio 2023, 13, 532–544. DOI:10.1002/2211-5463.13564

[38] W. Zhang, P. Xu, H. Zhang, Pectin in cancer therapy: A review. Tr. Food Sci. Technol. 2015, 44, 258–271. DOI:10.1016/J.TIFS.2015.04.001

[39] X. Liu, F. Wang, J. Wang, et al., Citrus flavonoid-pectin conjugates with enhanced emulsifying properties. Food Hydrocoll. 2025, 159, 110675. DOI:10.1016/j.foodhyd.2024.110675

[40] L. Wang, L. Zhao, F.I. Gong, et al., Modified citrus pectin inhibits breast cancer development in mice by targeting tumor-associated macrophage survival and polarization in hypoxic microenvironment. Acta Pharmacol. Sin. 2022, 43, 1556–1567. DOI:10.1038/s41401-021-00748-8

[41] X. Wang, Y. Yang, Y. An, et al., The mechanism of anticancer action and potential clinical use of kaempferol in the treatment of breast cancer. Biomed. Pharmacother. 2019, 117, 109086. DOI:10.1016/j.biopha.2019.109086

[42] A. V. P. Bobadilla, J. Arévalo, E. Sarro, et al., In vitro cell migration quantification method for scratch assays. J. R. Soc. Interface 2019, 16, 20180709. DOI:10.1098/rsif.2018.0709

[43] B. Brzozowska, M. Gałecki, A. Tartas, et al., Freeware tool for analysing numbers and sizes of cell colonies. Radiat. Environ. Biophys. 2019, 58, 109–117. DOI:10.1007/s00411-018-00772-z

